# Clinical-grade Plant-made nanomaterials: from Process Design to the construction of a Manufacturing facility

**DOI:** 10.1101/2025.10.14.682267

**Authors:** Denise Pivotto, Anthony Rosa, Aya Maged Elsheikh, Elisa Gecchele, Roberta Zampieri, Alessia Raneri, Valentina Garonzi, Linda Avesani

## Abstract

Plant-made nanomaterials are proteinaceous elements that are emerging as multi-purpose and versatile tools in the therapeutic landscape. In the context of autoimmune diseases, Tomato Bushy Stunt Virus (TBSV) has been previously explored as a platform for inducing immune tolerance by displaying disease-specific immunodominant peptides—offering a potential path toward disease remission.

In this study, we developed a dedicated facility and a GMP-compliant manufacturing process for producing TBSV-based nanoparticles engineered to display peptides relevant to specific autoimmune disorders.

Data collected from multiple non-consecutive pilot-scale production batches were used to build a simplified techno-economic model of the process.

The process is readily scalable and offers opportunities for further improvements, supporting the potential to meet market demands for early-stage therapeutic interventions in autoimmune diseases.

Additionally, an environmental, health, and safety (EHS) assessment of the process demonstrated a highly favorable environmental output index and minimal associated risks, reinforcing the platform’s sustainability.

These results support the viability of plant-based manufacturing for therapeutic nanomaterials and highlight TBSV’s potential as a novel platform for tolerance-inducing treatments in autoimmune diseases.

## Introduction

Plant Molecular Farming (PMF), which includes the production of recombinant proteins and nanomaterials using plant biotechnology, could provide a step change in improving health outcome, especially in developing countries and in pandemic contexts, thanks to its scalability, safety and rapidity.

The first cGMP manufacturing facility producing Plant-Made Pharmaceuticals (PMPs) was designed by Large Scale Biology Corporation (LSBC) in Owensboro, KY, USA (now Kentucky BioProcessing) and opened in the year 1999, using the plant-virus transient expression system Geneware® (Pogue et al., 2002, 2010). Ever since, PMF has progressed as a safe, easily scalable and cost-effective technology to produce biopharmaceuticals, with respect to other cell-based systems. Canadian former enterprise Medicago, now acquired by Aramis Biotechnology, was one of the pioneers in the industry: founded in 1999, its focus was the production of Virus-Like Particles (VLPs) as vaccines, reaching phase III clinical studies with their product Covifenz®, a vaccine candidate for the treatment of SARS-CoV2, before announcing closure in 2023.

Within this continuous framework and thanks to ongoing efforts in basic research, various emerging applications now rely on plant-made nanomaterials, primarily derived-from or inspired-by viral structures, as promising tools for addressing multiple human diseases. These materials represent a natural progression in the application of plant biotechnology to create innovative solutions for a range of human diseases.

Viruses are excellent examples of naturally occurring nanoparticles that can serve as ideal platforms for the development of new biomedical applications. Plant virus capsids are formed through the self-assembly of repeating protein subunits, therefore providing a high degree of multivalency. Plant viruses in general are non-enveloped structures, and they can assume a spherical/icosahedral or filamentous/tubular shape. Their repetitive structure and safety for humans make them a platform with peculiar characteristics featuring as adjuvants, active drugs and/or scaffolds for peptide and drugs display (Chung et al., 2020; Santoni et al., 2020).

While there is currently no clinically approved plant-based nanomedicine in the market, several are undergoing preclinical development while some systems are poised to enter translational development.

We recently demonstrated the potential of plant-made nanoparticles for preventing and curing autoimmune diseases such as Type 1 Diabetes (T1D) and Rheumatoid Arthritis (RA) (Zampieri et al., 2020). Indeed, the immense potential of nanomaterials as tolerogenic agents to be used in the context of diverse immunological diseases such as autoimmune diseases, allergies and transplants has been extensively demonstrated (Kusumoputro et al., 2023); however, plant viruses have never before been employed for such applications prior to our study.

For RA, we demonstrated that the use of plant-made nanoparticles was able to induce Treg cells by diverting the autoimmune response to a tolerogenic setting that reversed the clinical score of the disease with outcomes comparable to golden standard treatments based on immunosuppressants (Zampieri et al., 2020).

Given the promising preclinical evidence, we made a concerted effort to spin-out the project from an academic setting by establishing a GMP-grade manufacturing facility, enabling the production of nanoparticles under GMP conditions for use in human clinical studies.

The nanoparticles used are based on Tomato Bushy Stunt Virus (TBSV), a virus of the Tombusvirus family. TBSV is characterized by an icosahedral capsid of ~ 32 nm in diameter made up of 180 subunits of the coat protein (CP). TBSV Nano-Particles (NPs) can both encapsulate or expose small molecules and polypeptides on the surface (Grasso et al., 2013) and are neither toxic nor teratogenic (Blandino et al., 2015). When intravenously injected into mice, these NPs do not induce alterations of tissues/organs (Lico et al., 2016).

TBSV NPs manufacturing in plants relies on the use of transient viral expression in *N. benthamiana*, widely used as a bioreactor for the production of biopharmaceuticals, due to its versatility and susceptibility to plant pathogens, such as viruses commonly used as expression vectors (Bally et al., 2018). Besides acting as vectors that regulate gene expression, plant viruses can be genetically engineered to incorporate non-native peptides into their coat proteins. The modified viruses are then propagated in host plant leaves, which are subsequently harvested and processed for purification. (Pogue et al., 2002).

In this sense, once harvested after infection, the plant leaves can be processed to obtain a purified virus that exposes a target peptide: this whole NP represents the final API (Active Pharmaceutical Ingredient). Being a plant pathogen, the use of these NPs for therapeutical purposes is entirely safe for man, generally because of the absence of specific receptors for recognition and penetration into hosts’ cells (Nikitin et al., 2016).

Here, we describe the manufacturing process overseeing the production of NPs based on TBSV within a small-scale laboratory setting, meant to be GMP-compliant and designed to produce quantities sufficient for toxicological pre-clinical studies and Phase 1 clinical trials in humans. All projections presented throughout the article are based on a hypothetical API dosage of 2 mg per patient/year, aimed at achieving the desired clinical outcome of tolerance induction in RA patients.

## Materials and methods

### Host Plant Species Selection and Biomass Production

In the described experimental setup, *Nicotiana benthamiana* plants are cultivated indoors under controlled environmental conditions, maintaining a constant temperature of 24°C ± 3°C and relative humidity between 50% and 60%, with a photoperiod of 16 hours light and 8 hours dark. Seeds are initially sown and allowed to germinate for up to 10 days, after which the plantlets are moved into larger pots to accommodate growth and are redistributed across trays. Plants are irrigated every 2-3 days for a period of 18 days, until they reach optimal biomass. At this stage, the average fresh weight per plant is approximately 8.2 g. Five-week-old *N. benthamiana* plants are then ready for infection, initiating the manufacturing workflow, which is divided into upstream and downstream processing phases.

### Upstream Process

*N. benthamiana* plants used in the process are grown at a constant temperature of 24°C ± 3°C and a humidity of 50/60%, with a dark/light cycle of 16/8h.

The transient expression of TBSV vectors is achieved by *in vitro* transcription to produce TBSV.pLip infectious RNAs, of which 4 µg per plant are used for the manual infection of two leaves in *N. benthamiana* with the aid of an abrasive powder (Celite®). After 6 days, leaves displaying local and systemic infection symptoms are pooled and checked for recombinant virus RNA by reverse transcription polymerase chain reaction (RT-PCR). The collected leaves are homogenized in Phosphate Buffered Saline (PBS) 1X, composed of 151 mM NaCl, 8.4 mM Na_2_HPO_4_ × 12H_2_O, 1.86 mM NaH_2_PO_4_ × H_2_O, adjusted to pH 7.2), at a ratio of 1:10 (g LFW/mL buffer), resulting in a suspension containing virions, which is used in the next step.

40µl of sap are used to infect one leaf, for a total of two leaves per plant, with the same rubbing procedure described above; once again, the state of the infection is monitored by phenotypical assessment, then, 6 days post infection, the leaves are harvested and homogenized with PBS 1X to obtain another infectious suspension to proceed analogously with the infection of a third batch, for the third step of the process named tertiary infection. The plant biomass originated from this phase represents the starting material to be processed for final product purification. The necessary quality controls are performed during all phases, to ensure structural integrity of the virus and presence of the peptide of interest: these include RNA extraction for RT-PCR and sequencing, as well as Western blot (with a conformational anti-TBSV antibody) and Coomassie staining on a native agarose gel for TBSV NPs quantification.

Western blot analysis is performed using a 1% agarose, 38mM glycine gel electrophoresis in native conditions. For the electrophoretic run, samples are added a 6X loading dye (for 10ml: 6mL glycerol 100%, 1mL Tris-HCl 0.5M pH 6.8 and 18mg bromophenol blue). After inclusion of the loading dye, samples are directly loaded onto the agarose gel, which is then run at 100V for 45 minutes. Following electrophoresis, proteins are transferred onto a nitrocellulose membrane and the presence of TBSV is detected using 1:3000 anti-TBSV primary antibody (TBSV-CO – Prime Diagnostic) and horseradish peroxidase-conjugated anti-rabbit polyclonal antibody diluted 1:3000 (Clinisciences). The signal development is obtained on washed membranes by enhanced chemiluminescence (Amersham Biosciences, Amersham, UK) and the images acquired through ChemiDoc Imaging System (Biorad). For Coomassie staining, the gel is run in the same conditions and then stained with 30mL Quick Coomassie Stain (Clinisciences).

For RT-PCR, total RNA is extracted using protocols provided by the manufacturer of the TRIzol™ reagent (Invitrogen). Following RNA extraction, cDNA is synthesized, and PCR is carried out using primers TBSV_CP_For (TGCAACTGGTACGTTTGTCATATC) and TBSV_2_Back (AAGATCCAAGGACTCTGTGC). RNA extraction is the only part of the process that uses a small amount of hazardous chemicals, due to extraction with Trizol™ agent, such as chloroform and lithium chloride, but it is an essential step for quality control, and for which other commercially available extraction kits are under evaluation, to lower the environmental impact on production.

### Downstream Process

The downstream process begins with the collection of the infected leaves obtained from the tertiary cycle of infection. The biomass is first weighed to determine the fresh weight (FW) and then homogenized accordingly with 3 volumes of extraction buffer (plant-to-buffer ratio, weight in mg to volume in ml) using a steel blender, then filtered using four layers of Miracloth®. After this step, the homogenate undergoes a sedimentation process overnight at 4°C, to help precipitate debris and plant components. The precipitate is then consolidated with a first super-centrifuge step, after which the clarified supernatant is collected and undergoes an ultra-centrifugation round. Both centrifugation steps are performed with an Avanti JXN26 Centrifuge (Beckman Coulter). The pellet obtained after ultracentrifugation is resuspended in 0.7% physiological buffer solution (NaCl in Water for Injection), centrifuged once again and sterile filtered: this represents the final product, which then undergoes all quality control steps. For this process, at the end of both primary and secondary infection, the product undergoes RT-PCR, sequencing and a Western blot for confirmation respectively of RNA and coat protein identity. After the tertiary infection, the API is produced and the following controls are performed: Dynamic Light Scattering (DLS), Western blot, SDS-PAGE, DAS-ELISA for quantification and LAL test as described by the manufacturer to assess endotoxin content (Gen Script).

For SDS-PAGE analyses, TBSV NPs sample is supplemented with 0.5 volume of R buffer (for 10 ml: 3 ml of Tris-HCl 0.5 M pH 6.8, 2.4 ml glycerol 100%, 1.6 ml SDS 10%, 1 ml Bromophenol blue in TE, 300 µl EDTA 15 mM, 1.7 ml H_2_O; every 160 µl add 10 µl of 2-mercaptoethanol). After the addition of R buffer, samples are incubated at room temperature for 3 minutes and vortexed, then heated at 60 °C for 1 minute, vortexed again, and kept on ice until electrophoresis. TBSV NPs are then visualized using silver staining, following the manufacturer’s instructions (Pierce™ Silver Stain Kit, Thermo Scientific).

DAS-ELISA is used to determine TBSV concentration in solution. Briefly, 150 µl of sample diluted in diluent solution (0,2% BSA - PBS-Tween 0.05% pH 7.4) are distributed in ELISA Maxisorp plates (NUNC) previously coated with 150 µl of primary antibody TBSV-Co (Prime Diagnostic) diluted 1:1000 in carbonate buffer (Na_2_CO_3_ 1,59g/L, NaHCO_3_ 2,93g/L pH 9.6). The presence and quantity of TBSV NPs is revealed by adding 150µl secondary alkaline phosphatase conjugated antibody TBSV-AP (Prime Diagnostic) diluted 1:1000 in diluent solution and 100µl p-Nitrophenyl Phosphate (pNPP) substrate. The reaction is stopped using 100µl of 0.1M NaOH. Plate is read using a Tecan Infinite 200 PRO plate reader at 405 nm. To quantify TBSV NPs, the ELISA test includes a calibration curve ranging from 30.02 to 0.06 nanograms.

For DLS analysis, 0.4–0.5 mg/mL of TBSV NPs are analysed three times using a Zetasizer instrument. Each measurement represents the mean of 12 repetitions.

### Manufacturing facility

The facility described has been designed for the production of a GMP-compliant biopharmaceutical, with manufacturing volumes suitable to support Phase 1 Clinical Studies and toxicological studies. In all laboratory rooms air is filtered using HEPA filters, to avoid any source of contamination. After the laboratory rooms, an isolating door grants access to the production and downstream areas. All areas of production are subject to pressurization, with the three main chambers (growth chamber, infection chamber and downstream) presenting a higher pressure than the adjacent corridors, which in turn present a higher pressure than the outside laboratories. This allows for air to flow from production towards the outside environment, thus avoiding the entry of external contaminations. Prior to entering the infection and downstream chambers, the use of proper PPE (Personal Protective Equipment) and specific dressing is mandatory, as well as undressing is necessary prior to exiting. Irrigation of the plants is performed manually and to achieve optimal plant growth, the LED system has been designed to comprise three light channels: red (660nm), blue (450nm) and white (prevalently in the region between 500 and 600 nm). All LED have a certified degree of protection IP65, and an efficiency of 3.15µmoles/J. Each LED is 100cm long and four of them are meant to fit a 200 cm x 60 cm shelf, arranged for plants growth, and four shelves compose a rack, for a total capacity of 1024 plants in both the growth room and the infection room.

## Results

### Process and facility design

Plant-based biomanufacturing of therapeutic proteins represents a relatively novel platform with few commercial-scale facilities currently in operation. However, it provides several advantages, including linear production scalability, simplified upstream processes, shorter time to market, and the potential for reduced capital and operational expenditures.

This technology is at the core of Diamante SB, where *N. benthamiana* is used as a self-building, single-use, biodegradable bioreactor to produce a therapeutic nanomaterial named TBSV.pLip. The company facility has been carefully designed to accomplish this purpose, mindful of the active use of plant viruses in the production process. The equipment used for quality control and production is cGMP compliant. The laboratory rooms in the facility are equipped to perform all quality control analyses of the product. The current production process is controlled and monitored during working hours by working personnel. The high presence of manual operations during the production process, such as plant handling, plant-material extraction and centrifuge runs, at the moment do not allow for a more automated process, but remote monitoring is currently being implemented. Specific equipment like the −80°C freezer, used to stock material, and the general ventilation system are controlled remotely, allowing for tracking and fast intervention at any time.

An overview of the facility is shown in Figure 1.

**Figure 1:**
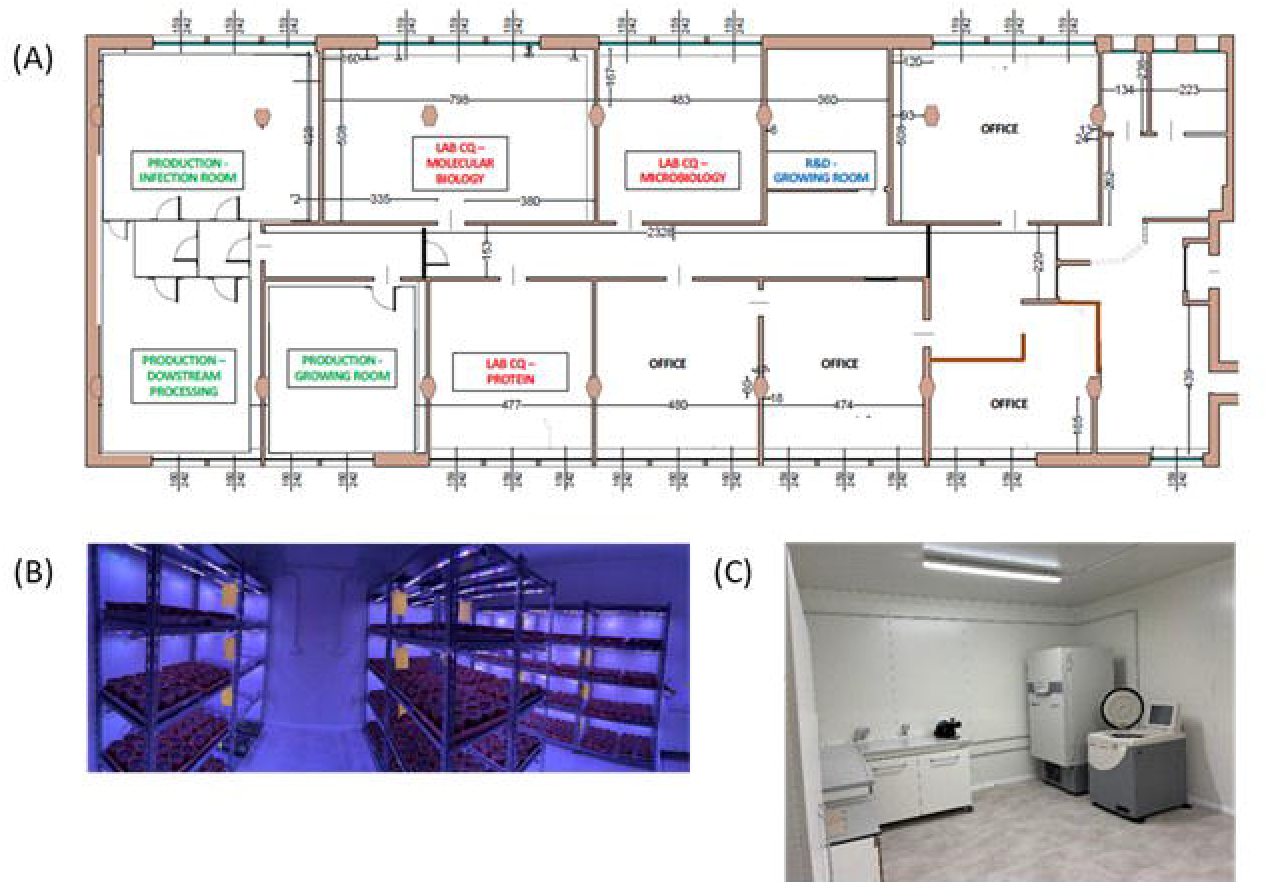
Overview of the facility. A) Facility floor plan. B) Production - growing room: together with the infection room, where the upstream process takes place. C) Production – downstream processing dedicated room.

### Manufacturing of TBSV.pLip NPs

TBSV.pLip NPs are produced using a process that can be summarized as described in Figure 2a. The upstream process is divided into three subsequential sections, beginning with the primary infection, as described previously (Lico et al., 2021). Once symptomatic leaves are harvested, they are extracted to produce SAP as starting material for a secondary infection. The same process is repeated with leaves collected from this second step, to continue with a tertiary infection, after which leaves are harvested and directly processed in the downstream phase: this latest phase ends with collection of TBSV.pLip. DAS-ELISA assay is used to correctly quantify the API: it has been estimated that the average total production yield for one batch of plants is 71mg/kg. This was calculated in relation to the plants used for all the production phases, as summarized in Figure 2b.

**Figure 2:**
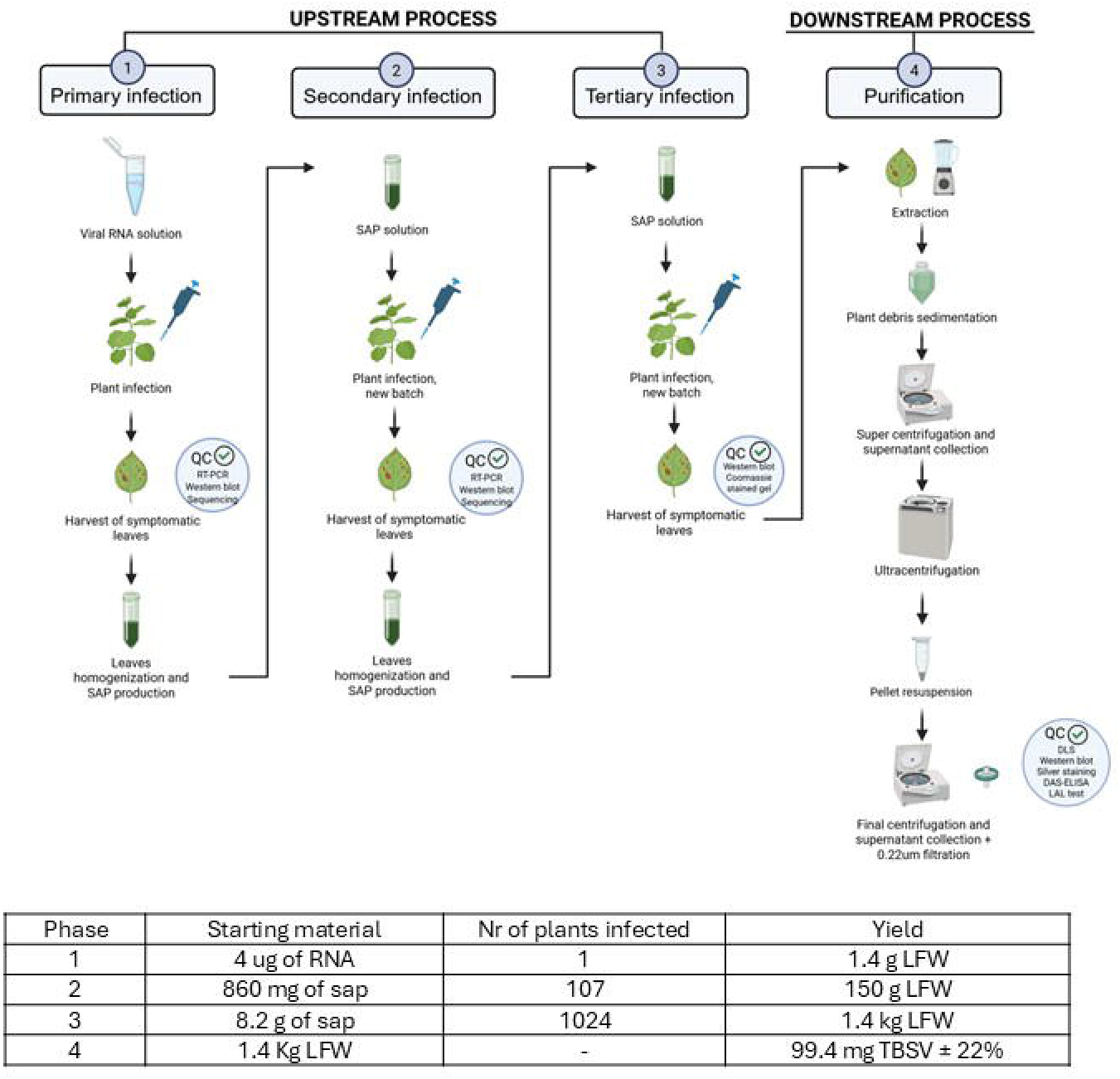
A) Process overview. The initial phase of the process, referred to as the upstream stage, is subdivided into three successive infection steps: primary, secondary, and tertiary. Following the tertiary infection, the product undergoes purification through the downstream processing phase. B) Starting material and process yields from the four phases described. LFW: leaf fresh weight.

For each stage of the process. *N. benthamiana* leaves are collected only when symptomatic: typical symptoms of virus presence in a plant include curling of the leaves and chlorosis (Figure 3). It is only these leaves that are processed to obtain the final product.

**Figure 3:**
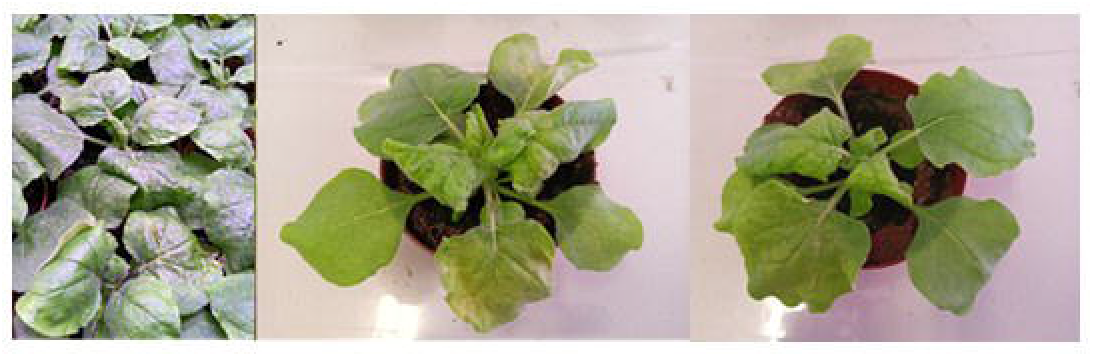
Plant infection. Example of symptomatic *N. benthamiana* plants infected with TBSV.

During each step of the process, the necessary quality controls are performed to ensure NPs integrity: RT-PCR, Western blot, Coomassie stained agarose gels and sequencing are part of the upstream phase, whereas in the downstream phase, the controls comprise DLS, SDS-PAGE, Western blot, DAS-ELISA and LAL Test (Figure 4).

**Figure 4:**
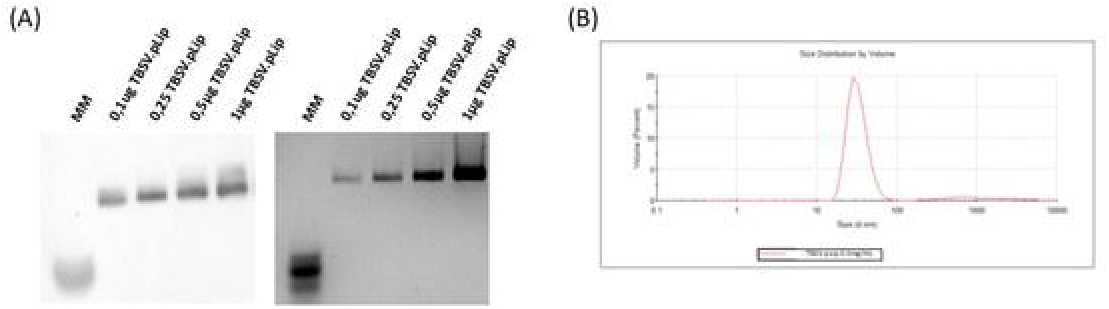
Analyticals overwiew. A) 1% agarose gel transferred to a nitrocellulose membrane and blottet against an anti-TBSV antibody and 1% agarose gel colored with Coomassie Quick Stain show a calibration curve with increasing quantities of TBSV.pLip; B) DLS profile of TBSV.pLip sample concentrated 0.5mg/mL, indicating distribution by volume: the peak indicates that most of the population is in the range of 30nm, which is the molecular size of the virus

### Economic Analysis of TBSV Production Costs

Annually, the production system in the current facility allows for cultivation and processing of 13 full batches, each comprising 1,024 plants. Due to the capacity of the available ultracentrifuge, which is the primary bottleneck in downstream processing, a maximum of 256 plants can be processed per day. As a result, each full batch is processed over four consecutive days, in sub-batches of 256 plants per day. The processing of a single sub-batch yields approximately 24.9 mg of TBSV. Therefore, the total yield per full batch is: 99.4 mg of TBSV.pLip. Given the production of 13 batches per year, the estimated annual yield is: 1.3 grams of TBSV.pLip. To obtain 1 gram of TBSV.pLip, 10 full batches are required.

A comprehensive economic analysis was conducted to assess the operational expenditure (OPEX) associated with the production of 1 gram of TBSV and it is reported in Table 1 with the corresponding graphical illustration in Figure 5. This analysis is based on current small-scale batch production data and includes both direct and indirect costs, while capital expenditures (CAPEX) such as facility depreciation, equipment amortization, and long-term investments are excluded from the present assessment.

**Table 1:** Breakdown of the process cost per category, including direct and indirect costs. All cost values reported here reflect the aggregate expenses incurred to produce 1 gram of TBSV, inclusive of upstream (US) and downstream (DS) operations, as well as quality control (QC) and managerial oversight.

| Component | Percentage | Component | Percentage |
| --- | --- | --- | --- |
| Direct Costs |  |  |  |
| Raw Materials & Consumables | 1% | Materials & Consumables | 5% |
| QC Materials & Consumables | 4% |  |  |
| Production Labor Cost | 14% | Labor Cost | 40% |
| Production Manager Cost | 11% |  |  |
| QC Labor Cost | 6% |  |  |
| QC Manager Cost | 9% |  |  |
| Indirect Costs |  |  |  |
| Electricity | 5% | Utilities | 8% |
| Water | 1% |  |  |
| Heating | 2% |  |  |
| Rent | 40% | Facility-Related Costs | 47% |
| Building Maintenance | 3% |  |  |
| Cleaning Services | 5% |  |  |
| Total | 100% | Total | 100% |

**Figure 5:**
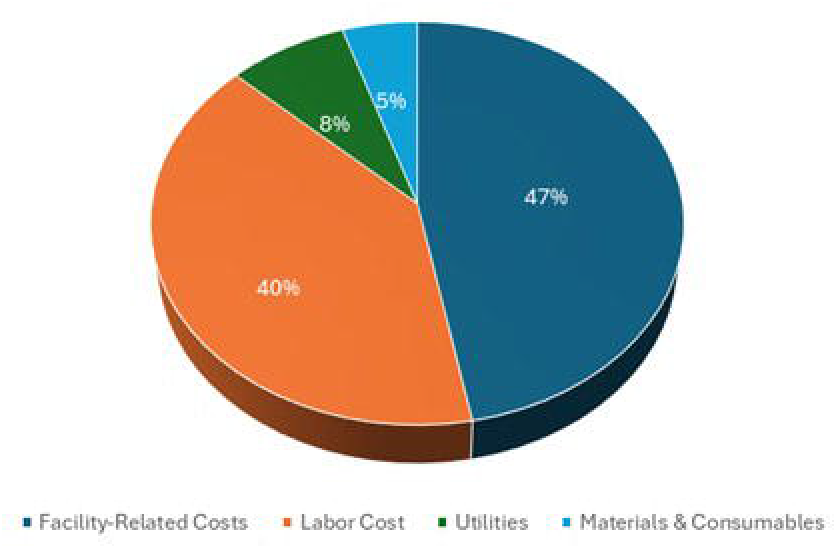
Breakdown of the percentage contribution of each cost category to the total production cost.

The results highlight that labor-related costs constitute the most significant portion of the annual operational direct cost, representing 40% of total OPEX. This includes both production labor (14%), QC labor (6%), and their respective managerial roles (11% for production management, 9% for QC management). These findings reflect the labor-intensive nature of the process, especially under small-scale conditions requiring high oversight and manual handling. Facility-related expenses emerged as the second major cost component, contributing 47% of total OPEX. The largest element within this category is facility rent, which alone accounts for 40%, followed by cleaning services (5%) and building maintenance (3%). These are fixed costs that remain constant regardless of output volume, thereby exerting a strong influence on cost per unit at limited production scales. Utilities, including electricity (5%), water (1%), and heating (2%), together contribute 8% of total OPEX. These costs are primarily associated with machinery operation and climate control, particularly during downstream processing steps. In contrast, materials and consumables represent only 5% of the total OPEX. This includes raw materials (1%) and quality control reagents and consumables (4%). Despite being essential to the production process, these inputs have minimal economic impact compared to labor and infrastructure costs.

### Upstream vs. Downstream Cost Distribution

In the upstream phase, including cultivation, infiltration, and incubation, the dominant costs are labor-related. Production labor and its associated management constitute a combined 25% of the total OPEX, while QC labor and materials used during in-process checks contribute an additional 5%. The use of raw materials in upstream operations accounts for a modest 1% of the total operational budget. In the downstream phase, comprising tissue harvesting, clarification, ultracentrifugation, and product release testing, labor remains the primary cost driver. QC labor and QC management represent a combined 15%, reflecting the intensive resource demands of analytical validation and batch release procedures. Production labor associated with downstream extraction adds another 3%–5%, while QC materials and consumables contribute 4%. The cost of raw materials used in this phase is negligible (<1%).

### Spatial Productivity and Vertical Farming Integration in Plant-Based TBSV Production

An essential parameter for evaluating the scalability of plant-based production platforms is spatial productivity, commonly expressed as yield per unit area. In the current system, the TBSV.pLip yield has been estimated at 6.2 mg per square meter of cultivation area per batch. Assuming 12 months of active production per year, this corresponds to an annual yield of approximately 80.6 mg per square meter per year. Assuming an annual therapeutic dose of 2 mg of TBSV.pLip per patient (a dosage that remains to be validated for clinical studies), this production rate allows one square meter of cultivation space to support treatment for approximately 40 patients per year. To meet a therapeutic demand of 5,000 patients per year, an estimated 125 square meters of cultivation area would be required. This requirement increases proportionally with treatment demand, reaching 1,250 square meters and 25,000 square meters for 50,000 and 1,000,000 patients, respectively (Table 2). To optimize land use efficiency and reduce the physical footprint of the production facility, a vertical farming system should be adopted (Figure 6). By incorporating five levels of vertically stacked shelving, the required floor area is effectively reduced by a factor of five. Accordingly, the floor area needed to treat 5,000, 50,000, and 1,000,000 patients per year is reduced to 25 m^2^, 250 m^2^, and 5,000 m^2^, respectively (Table 2).

**Table 2:**
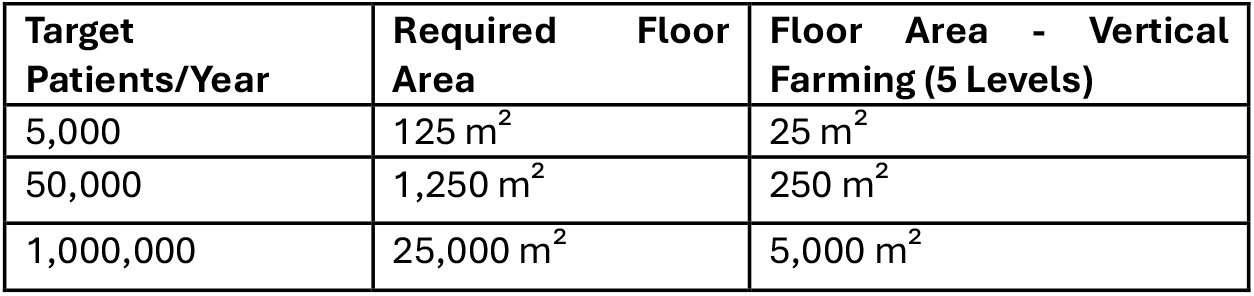
scalability of plant-based NPs production by using vertical farming.

**Figure 6:**
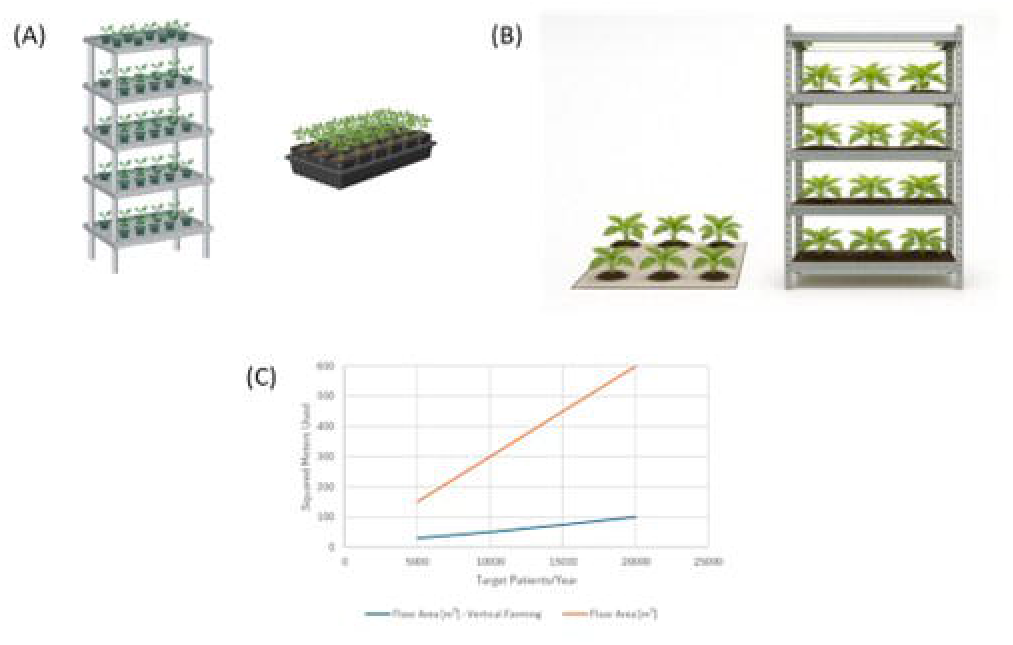
(A) (B) The advantage of vertical farming versus traditional plant cultivation. The use of shelves allows to take advantage of the entire room area, extending cultivation in vertical layers and optimizing the space (C).

This spatial modularity offers a clear advantage for scalable models, enabling increased productivity per unit footprint and facilitating expansion without the need for large horizontal land allocations. When combined with the inherent flexibility of plant-based systems, vertical farming provides a strategically efficient pathway for transitioning toward larger-scale, economically sustainable production of biopharmaceuticals such as TBSV, as illustrated in Figure 6.

### Materials Hazardness and Environmental Impact

The plant-based transient expression system used in this study is inherently more sustainable than traditional production platforms, largely due to the biodegradable nature of plant biomass and the low-impact cultivation requirements of *Nicotiana benthamiana*. Plants serve as renewable, carbon-sequestering bioreactors, and their post-harvest waste can often be composted or disposed of with minimal environmental burden (Buyel, 2019). Nonetheless, like many biotechnological processes, our workflow includes the use of a limited number of hazardous chemicals, such as Trizol®, chloroform, hydrochloric acid, and lithium chloride. Importantly, these substances are utilized in minute quantities—typically in microliters per batch—and their contribution to the overall environmental footprint is negligible. Strict adherence to laboratory safety and waste management protocols further mitigates their risk.

One potential concern unique to our system is the use of the TBSV as a vector. While TBSV is a plant-specific virus with no known risk to humans or animals, its presence in biomass waste could raise biosafety considerations, particularly in open-field or large-scale settings. We recognize this challenge and are currently investigating strategies to inactivate the virus post-harvest, ensuring that the final biomass is free of viable viral particles before disposal or downstream use. In summary, the combination of (i) low-volume reagent usage, (ii) elimination of persistent organic solvents, (iii) the biodegradable nature of plant material, and (iv) ongoing viral inactivation efforts, reinforces the environmental advantages of our production method.

## Discussion

Despite the recent closure of Medicago following the withdrawal of investment by its parent company, the global landscape of plant-based biotechnology companies remains dynamic. Notable examples include Protalix Therapeutics (Israel), with two marketed products—Elelyso® and Elfabrio®; BioApp (South Korea), developer of the HERBAVAC™ pig vaccine; KBio (USA), focused on various recombinant proteins; Eleva (Germany), advancing a moss-derived therapeutic for Fabry disease; InVitria (USA), producing technical-grade proteins; ORF Genetics (Iceland), supplying growth factors and cytokines for cosmetics and research; Agrenvec (Spain) and Baiya Phytopharm (Thailand), both targeting applications in cultivated meat and cosmetics; and CapeBio (South Africa), developing medical diagnostic solutions.

In this vibrant and dynamic scenario, Diamante SB, founded in 2016 as a spin-off of the University of Verona with support from European public funding programs such as Bio-based innovation for sustainable goods and services with the project PharmaFactory and the EIC accelerator, has successfully transitioned from an academic setting to an industrial reality. The company is now focusing on the development of a platform for autoimmune disease treatment with the first therapeutic application in Rheumatoid Arthritis. Here we reported a basic techno-economic analysis from laboratory data collected in the first year of work in the new lab-setting meant to obtain GMP certification disclosing the production process of the therapeutic TBSV.pLip meant for RA treatment.

The facility described annually produces 1.3 g of TBSV.pLip. The host used is *Nicotiana benthamiana*, selected for its productivity and since host of TBSV infection. Furthermore, *N. benthamiana* is familiar to EMA and FDA thus facilitating its acceptance in regulation-compliant manufacturing. (Streatfield and Howard, 2003; McCormick et al., 2008; Bendandi et al., 2010; Tusé, 2011; Plant Viral Vectors for Delivery by Agrobacterium, 2013)

The API is manufactured in *N. benthamiana* using TBSV genome as a vector of expression and exploiting the TBSV Coat Protein for the display of the peptide Liprin (pLip) on the external surface of the capsid.

Each batch of 1024 plants, housed in vertically stacked racks under high-efficiency LED lights, grows over the course of 6 days and yields a total of 1.4 kilograms (fresh weight) of biomass.

13 batches are seeded and grown annually, with one batch reaching harvest every 28 days. Expression rate of 355 mg of TBSV.pLip per kilogram of biomass (fresh weight) and a downstream recovery of 20% give a yield of 71 milligram of TBSV.pLip per kilogram of harvested biomass.

The economic feasibility of TBSV production at small scale was conducted by analyzing direct operating expenditures (OPEX), which include labor, materials, and utilities in both upstream and downstream processes. Capital expenditures (CAPEX) were excluded to focus on recurrent, process-dependent costs. As detailed in the Results section, labor and facility-related costs dominated the cost structure, accounting for 40% and 47% of total OPEX, respectively, while materials and consumables contributed only 5%.

Based on the annual yield of approximately 1.3 grams, the direct OPEX per mg of TBSV was estimated at €27.61. This cost reflects the fixed-intensive nature of the process and highlights the potential for improvement through economies of scale. Increasing batch numbers or optimizing facility use would reduce per-unit costs significantly.

Moreover, the labor-heavy profile suggests that selective automation, especially in routine handling and monitoring, could reduce manual workload, improve consistency, and support future scale-up. Together, these strategies represent key opportunities to enhance the economic sustainability of TBSV-based production, a platform that may be explored for diverse tolerance induction applications simply changing the peptide displayed by VNPs.

Finally, it is important to note that in addition to the conventional cost of goods, Life Cycle Assessment (LCA) accounting for the costs of the environmental footprint of manufacturing should be addressed, even in the field of biopharmaceuticals.

When comparing our process with standard methods used for the chemical synthesis of peptides, the core therapeutic agent, the footprint of the plant-based upstream process is negligible when compared to the volumes of organic solvents used in traditional solid-phase peptide synthesis (SPPS). In contrast, SPPS is well-documented for its environmental burden, primarily due to its reliance on toxic, non-biodegradable solvents (e.g., dimethylformamide, dichloromethane) and repetitive washing steps, which generate significant chemical waste. The cumulative impact of these solvents not only raises health and safety concerns but also requires energy-intensive waste treatment and solvent recovery systems.

This approach represents a meaningful step toward greener, safer, and more scalable biomanufacturing when compared with traditional peptide synthesis platforms and it could be used as a platform for the production of nanomaterials meant for tolerance induction in the framework of autoimmune diseases.

## Acknowledgments

This work has been co-funded by the EIC 2024 Accelerator program, specifically through the “Diamante” project under grant agreement number 101189019.

Authors acknowledge CPT (Centro Piattaforme Tecnologiche) of the University of Verona for the access and support to the DLS equipment.

Part of the work has been supported by ‘Finanziamento dell’Unione Europea-Next Generation EU, Missione 4, Componente 1 CUP B53D23024740001’.

